# Global terrestrial nitrogen uptake and nitrogen use efficiency

**DOI:** 10.1101/2022.11.01.514661

**Authors:** Yunke Peng, Iain Colin Prentice, Keith J. Bloomfield, Matteo Campioli, Zhiwen Guo, Yuanfeng Sun, Di Tian, Xiangping Wang, Sara Vicca, Benjamin D. Stocker

## Abstract

Plant biomass production (BP), nitrogen uptake (*N*_up_) and their ratio, nitrogen use efficiency (NUE), must be quantified to understand how nitrogen (N) cycling constrains terrestrial carbon (C) uptake. But the controls of key plant processes determining *N*_up_ and NUE, including BP, C and N allocation, tissue C:N ratios and N resorption efficiency (NRE), remain poorly known. We compiled measurements from 804 forest and grassland sites and derived regression models for each of these processes with growth temperature, vapour pressure deficit, stand age, soil C:N ratio, fAPAR (remotely sensed fraction of photosynthetically active radiation absorbed by green vegetation) and growing-season average daily incident photosynthetic photon flux density (gPPFD) (effectively the seasonal concentration of light availability, which increases polewards) as predictors. An empirical model for leaf N was based on optimal photosynthetic capacity (a function of gPPFD and climate) and observed leaf mass-per-area. The models were used to produce global maps of *N*_up_ and NUE. Global BP was estimated as 72 Pg C/yr; *N*_up_ as 950 Tg N/yr; and NUE as 76 gC/gN. Forest BP was found to increase with growth temperature and fAPAR and to decrease with stand age, soil C:N ratio and gPPFD. Forest NUE is controlled primarily by climate through its effect on C allocation – especially to leaves, being richer in N than other tissues. NUE is greater in colder climates, where N is less readily available, because belowground allocation is increased. NUE is also greater in drier climates because leaf allocation is reduced. NRE is enhanced (further promoting NUE) in both cold and dry climates. These findings can provide observationally based benchmarks for model representations of C–N cycle coupling. State-of-the-art vegetation models in the TRENDY ensemble showed variable performance against these benchmarks, and models including coupled C–N cycling produced relatively poor simulations of *N*_up_ and NUE.

## INTRODUCTION

Although the main fluxes in the terrestrial carbon (C) cycle are relatively well quantified, large uncertainty surrounds many aspects of the nitrogen (N) cycle (Davies-Barnard et al., 2020; Zaehle et al., 2014). The two are inextricably linked (Cleveland et al., 2013; Hungate et al., 2003) due to the N requirements for plant growth (Norby et al., 2010), with N availability influencing the relationships among photosynthesis and growth (LeBauer & Treseder, 2008; Liang et al., 2020; Vicca et al., 2012), biomass production (BP) and allocation (Fay et al., 2015; Poorter et al., 2012; Terrer et al., 2019), and rhizodeposition (Henneron et al., 2020; Perkowski et al., 2021). Yet, recent ecosystem models have reported global plant N uptake (*N*_up_) ranging widely, from 465 to 1197 Tg N yr^−1^ (Cleveland *et al*., 2013; Oleson *et al*., 2010; Smith *et al*., 2014; Goll *et al*., 2017b; Lawrence *et al*., 2019; Wiltshire *et al*., 2021). Although N cycling processes are expected to constrain the response of terrestrial C uptake to rising CO_2_ (Hungate et al., 2003), the assumptions about these processes made in current models vary considerably and global compilations of site-level C and N cycling remain underutilized for the their evaluation (see e.g. Zaehle *et al*., 2014; Fowler *et al*., 2015). As a consequence, process-based models make widely divergent predictions of the extent of N limitation to global C uptake in scenarios of future CO_2_ and climate change (Zaehle *et al*., 2014; Stocker *et al*., 2016; Arora *et al*., 2020).

Meanwhile, observational studies have generated a substantial body of ecosystem-level observations relevant to N cycling that has not previously been used in model development or evaluation. Here we use such data, derived from multiple sources, to fit and upscale statistical models of key processes contributing to the terrestrial N cycle, with a view to providing new benchmarks to test (and potentially, better constrain) process-based models.

The starting point for our analysis is BP, which is distinct from net primary production (NPP). NPP is defined as gross primary production (total photosynthetic carbon fixation) minus plant respiration, while BP is the annual C actually used for the growth of leaves (BP_leaf_), wood (BP_wood_) and roots (BP_root_) (Collalti et al., 2020; Collalti & Prentice, 2019; Vicca et al., 2012). NPP includes the production of non-structural C compounds, including labile carbohydrates, volatile organic compounds (VOC) and root exudates (Collalti et al., 2020; Vicca et al., 2012) that do not form part of BP. However, although BP is readily available from field measurements (albeit with uncertainties – especially about the below-ground contribution, and the variable contribution of non-structural carbohydrates to measured BP), NPP generally is not. Our particular focus is then on *N*_up_ and on nitrogen use efficiency (NUE), which is the ratio of BP to *N*_up_. To estimate *N*_up_ and NUE, we analyse the environmental dependencies of the various components contributing to determining N cycling rates including BP, biomass allocation, tissue C:N ratios and N resorption efficiency (NRE).

BP has been found to increase with growth temperature (Baig et al., 2015) and soil nutrient availability (LeBauer & Treseder, 2008). A decline in forest BP with stand age is also well documented (Ryan et al., 2004; Xia et al., 2019). Few attempts have been made to describe global variations of *N*_up_ and NUE (but see Cleveland et al., 2013; Wang et al., 2018b). NUE has been indicated to increase as N supply becomes more limiting (Finzi et al., 2007; Harrington et al., 2001), and to be reduced at increased soil nitrogen-to-phosphorus (N:P) ratios (Gill & Finzi, 2016) or after N fertilization (Davies-Barnard et al., 2020). Variations in biomass distribution between different organs and their distinct C:N stoichiometry (Ma et al., 2021; Tian et al., 2019; Zhang et al., 2020), and variations in NRE (Deng et al., 2018; Du et al., 2020), must influence NUE, but there is limited knowledge of how these factors change along environmental gradients and of their importance in affecting *N*_up_ and NUE variations among sites with different climatic and edaphic conditions. Leaf stoichiometry varies greatly between species (Tian et al., 2019) but also shows systematic relationships with climate (Reich et al., 2007) and soils (Maire et al., 2015). Variations in mass-based foliar N content (*N*_mass_, mg g^−1^) have been interpreted as reflecting plant nutritional status (e.g. Penuelas et al., 2020) but *N*_mass_ depends in part on leaf mass per unit area (LMA) and in part on the amount of N invested in Rubisco, the key enzyme determining photosynthetic capacity (Dong et al., 2017; Luo et al., 2021). Photosynthetic capacity can be quantified by the maximum rate of carboxylation (*V*_cmax_). When standardized to 25°C, (*V*_cmax25_) this rate is related to the amount of Rubisco in leaves, and therefore to the amount of N per unit leaf area (Harrison et al., 2009). A substantial proportion of *V*_cmax25_ variation can be predicted by climate (Peng et al., 2021; Smith et al., 2019) and the same is true for LMA (Dong et al., 2022a; Wang et al., 2021), implying that foliar N content is at least partly controlled by climate. NRE has been shown to be negatively related to temperature and humidity. Since rates of N cycling are enhanced in warmer and wetter environments, N supply from resorption becomes relatively less important under these conditions: as temperature and humidity increase, N cycling rates shifts from the (more conservative) resorption pathway to the mineralization pathway (Deng et al., 2018; Du et al., 2020). Taking these various findings together, it seems likely that climate-driven demand for N – controlled by the relationships between climate and growth, allocation, tissue C:N stoichiometry and N resorption – as well as soil N supply – jointly determine the amount of N required to construct new biomass, and the efficiency with which the uptake of N is translated into plant growth.

BP has been simulated by using satellite products (Cleveland et al., 2013; Zhao & Running, 2010), or with models entirely driven by climate (Goll et al., 2017; Lienert & Joos, 2018; Mauritsen et al., 2019; Meiyappan et al., 2015). However, variations in C allocation with climate and soil nutrient availability, and their implication for N uptake, have typically been underestimated in models (Medlyn et al., 2015; Zaehle et al., 2014). As a step towards remedying this situation, we compiled a new global dataset of the key components determining N cycling rates in terrestrial ecosystems (BP, allocation, plant C:N stoichiometry and NRE) and associated environmental drivers and vegetation characteristics (climate, vegetation cover, stand age and soil C:N ratio), and analysed their inter-relationships using statistical methods (Fig. S1). We upscaled and combined the resulting statistical models to produce global maps of BP, *N*_up_ and NUE. We also made a first assessment of the ability of process-based terrestrial C and C-N cycle models from the TRENDY ensemble to represent the environmental responses of BP, *N*_up_ and NUE as shown in our analysis.

## METHODS

The analysis was conducted in six stages: (1) Compilation of a dataset of previously published stand-scale measurements for forest and grassland sites; (2) Global compilation of data to be used as predictors in statistical models; (3) Fitting statistical models at the stand scale; (4) Global application of the models, in order to estimate global terrestrial C and N uptake; (5) Analysis of the factors contributing to modelled *N*_up_ and NUE; and (6) Comparison of the fitted statistical models with simulations by state-of-the-art global vegetation models.

### Stand-scale datasets of plant and leaf traits

Our plant-trait dataset comprises measurements of total BP (g C m^−2^ yr^-1^) and the BP of leaves, wood and roots (Anderson-Teixeira et al., 2018; Campioli et al., 2015; Luyssaert et al., 2007; Malhi et al., 2011, 2017; Tian et al., 2019; Vicca et al., 2012; Wang & Zhao, 2022). 87% of the data are from forests, 13% from grasslands. 17%, 62% and 21% of the data are from tropical (0–22.5°), temperate (22.5°– 50°) and high-latitude (>50°) regions respectively. BP_leaf_, BP_wood_ and BP_root_ represent leaf, wood and root production, with BP_wood_ equal to zero in grasslands. Total aboveground BP (ABP) is the sum of BP_leaf_ and BP_wood_. Belowground BP (BBP) is equal to BP_root_. For a subset of the sites, we also obtained data on leaf C:N ratios, which were used to calculate the leaf N flux (BP_leaf_ divided by the leaf C:N ratio, g N m^−2^ yr^−1^).

We assembled an additional leaf-trait dataset including *N*_area_, *V*_cmax25_ and leaf mass per area (LMA), comprising 350 sites and 2424 species in natural (unfertilized) vegetation (Atkin et al., 2015; Bahar et al., 2017; Bloomfield et al., 2019; Cernusak et al., 2011; Domingues et al., 2010; Dong et al., 2017; Ferreira Domingues et al., 2015; Maire et al., 2015; Meir et al., 2017; Walker et al., 2014; Wang et al., 2018a; Xu et al., 2021). Nitrogen resorption efficiency (NRE) data were obtained from published sources at 210 sites (Deng et al., 2018; Du et al., 2020).

For comparison with *N*_up_, we used a forest net mineralization rate (*N*_min_, g N m^−2^ yr^-1^) dataset from 225 samples at 84 sites (Gill & Finzi, 2016). Net mineralization is the microbial release of inorganic N into the soil. This comparison rests on the assumption that annual ecosystem N gains and losses are small compared to mineralization and uptake. *N*_min_ may underestimate *N*_up_, as a significant contribution to *N*_up_ can be via organic forms of N (Liu et al., 2017; Näsholm et al., 2009). However, since direct measurements of *N*_up_ are not possible (and ecosystem *N*_up_ estimates are commonly derived from the same component fluxes and stoichiometry data that were used in our model development), *N*_min_ was considered here as an acceptable independent point of comparison for modelled *N*_up_ (Gill & Finzi, 2016).

### Global datasets of predictors

Maps on a half-degree global grid were developed for *V*_cmax25_ (μmol m^−2^ s^−1^), mean daytime air temperature (*T*_g_,°C), vapour pressure deficit (*D*, kPa), incident photosynthetic photon flux density averaged over the growing season (gPPFD, μmol m^−2^ s^−1^), the fraction of absorbed photosynthetically active radiation (fAPAR, unitless: a remotely sensed measure of green vegetation cover), stand age (years), soil C:N ratio (g g^−1^) and leaf mass per area (LMA, g m^−2^; Fig. S2). Variables were three-dimensionally (latitude, longitude, elevation) interpolated to site locations using Geographically Weighted Regression (GWR) in the ‘*spgwr*’ package in R to obtain plot-level predictors for empirical model fitting.

The global map of *V*_cmax25_ was obtained using a climatically driven model for *V*_cmax_, based on eco-evolutionary optimality principles (Peng et al., 2021; Prentice et al., 2014; Stocker et al., 2020; Wang et al., 2017). Global patterns of *V*_cmax_ predicted by this model have been shown to compare well to independent estimates derived from remotely sensed chlorophyll measurements (Dong et al., 2022b). *V*_cmax_ was predicted using Eq. C4 in Stocker *et al*. (2020), from atmospheric pressure, CO_2_, gPPFD and other daily climate forcing data (relative humidity, precipitation and average daily temperature) derived from WATCH Forcing Data ERA-Interim (WFDEI: Weedon et al., 2014). Values were converted to a standard temperature of 25°C (*V*_cmax25_) using the Arrhenius equation, with activation energy from Bernacchi et al. (2001). This converts *V*_cmax_ predicted by the model, which applies to growth temperature and therefore reflects optimality under natural field conditions, to a quantity (*V*_cmax25_) assumed proportional to Rubisco amount – and thus to the metabolic component of leaf N (Dong et al., 2017, 2022a). Estimated *V*_cmax25_ was averaged over 1982–2011, using the maximum daily *V*_cmax25_ value for each year.

Monthly average values of mean daily maximum (*T*_max_, °C) and minimum (*T*_min_, °C) temperature were obtained from Climate Research Unit data (CRU TS 4.0) (Harris et al., 2014) for the period 1980– 2016. *T*_g_ was estimated monthly by approximating the diurnal temperature cycle with a sine curve, where daylight hours are determined by month and latitude:

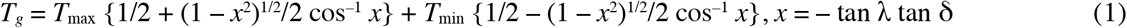

where *λ* is latitude and *δ* is the monthly average solar declination (Jones, 2013). Monthly values of *T*_g_ were averaged from 1980 to 2016, over the thermal growing season, i.e., months with *T*_g_ > 0 °C. Vapour pressure deficit (*D*) was estimated using gridded actual vapour pressure (*e*_a_, hPa) from

CRU, for the same period and resolution as *T*_g_, using GWR:

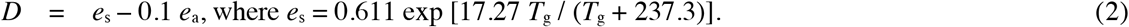

Monthly values of *D* were averaged from 1980 to 2016 over the thermal growing season.

Incident solar radiation data were obtained from WFDEI for the same period and resolution as *D* and *T*_g_. Solar radiation (W m^−2^) was converted to gPPFD assuming an energy-flux ratio of 4.6 μmol J^−1^ and a photosynthetically active fraction of 0.5. Monthly values were averaged from 1980 to 2016 over the thermal growing season.

Fraction of absorbed photosynthetic radiation (fAPAR) data were derived from Advanced Very High Resolution Radiometer (AVHRR) Normalized Difference Vegetation Index third generation (NDVI3g) map products for the period 1982–2011 (Pinzon & Tucker, 2014). Mean stand age was derived from Poulter et al. (2018), calculated as the mean age across four plant functional types (PFTs) and 15 age classes (from 0–10 to 140–150 years), weighted by their respective fractional area coverage within each grid cell. Missing values for stand ages were filled by the average of the local continent. Soil C:N ratio (Batjes, 2015) was processed by calculating the layer depth-weighted mean across the top 2-3 layers (20–60 cm). Global soil C:N maps were then aggregated from 1/120 to 1/2 degrees and spatially interpolated to fill the 8% of land area with missing C:N values based on the *k*-nearest-neighbour (KNN) method, using longitude and latitude as predictors and an optimized *k* = 7. LMA was also gap-filled by the KNN method (as predicted by latitude, *T*_g_, gPPFD and the ratio of actual evapotranspiration to potential evapotranspiration, optimal *k* = 9), filling 71% of the land area.

### Empirical models

Statistical models for BP, the allocation of BP to separate tissues, tissue C:N ratios and NRE were developed based on data from globally distributed forest and grassland sites (Fig. S3). We used interpolated (rather than directly measured) values to avoid the ∼86% reduction of sample size which would have occurred otherwise. Additional analyses, to test the validity of this choice, were carried out using directly measured values only.

For forests, a dataset of measured BP (*n* = 514; Fig. S3), ABP (*n* = 709), BP_leaf_ (*n* = 636) and site-level predictors interpolated from map products was used to fit statistical models. BP and its allocation have previously been modelled as functions of stand age (Campioli et al., 2015), soil fertility (Vicca et al., 2012) and climate (Collalti et al., 2020). Accordingly, we initially selected the following six variables for predicting BP, ABP/BP and BP_leaf_ /ABP in forest: stand age, soil C:N ratio, fAPAR, *T*_g_, gPPFD and vapour pressure deficit. Soil C:N provides an inverse indicator of soil N availability (Vicca et al., 2018). fAPAR represents an inverse measure of environmental stress – for example, due to a short growing season or to low light, water or nutrient availability. gPPFD is in effect a measure of the seasonal concentration of light availability and, unlike total growing-season PPFD, increases towards the poles. (The selection of predictors was nearly unchanged if total PPFD was used instead of gPPFD to predict BP, except that total PPFD was finally not selected, and *D* was newly included.)

Model selection (Table S1) was performed by forward stepwise regression, adding the variable producing the largest increase in *R*^2^ at each step. We required all variables included in the final model to have regression coefficients significantly different from zero (assessed by the *t*-statistic). Variables were added one-by-one until this criterion was no longer met. All ratios constituting response variables in the statistical models (ABP/BP, BP_leaf_/ABP, NRE) were logit-transformed, because these ratios range from 0 to 1; thus, values after transformation are continuous and unbounded, consistent with the assumptions of ordinary linear regression. For the same reason, stand age, soil C:N ratio, gPPFD and *D* were log-transformed. In one case – logit (BP_leaf_/ABP) – the model selection procedure failed, as age was identified as the first predictor but became non-significant as more variables were added. In this case, we repeated the model selection with stand age removed.

Since multiple individual trees may have been measured at each site, a mixed effects model was applied with site as the grouping variable for random offsets. Variance inflation factors (VIF) were calculated to test for multicollinearity in the BP and allocation models (Fig. S4). *T*_g_ and *D* were found to cause multicollinearity in BP and ABP/BP models (VIF > 10), so in these two models we included *T*_g_ but not *D*.

Following Dong *et al*. (2017), leaf *N*_area_ can be well approximated as the sum of a bulk leaf tissue component proportional to LMA, and a metabolic component proportional to *V*_cmax25_:

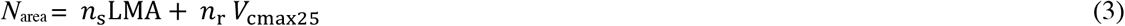

where *n*_s_ and *n*_r_ are empirical coefficients. It follows that:

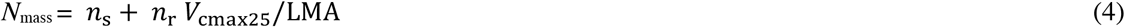

We fitted this *N*_mass_ model using parallel observations of *N*_mass_, *V*_cmax25_ and LMA, for 350 global sites and 2424 species. Here site and species were treated as grouping variables for random offsets using a linear mixed effects model with a crossed random design. To estimate leaf C:N ratio (*C*_mass_/*N*_mass_), leaf *C*_mass_ was assumed globally constant at the median value (0.47 g g^-1^) of the relevant subset of our leaf-trait dataset (*n* = 79 sites, 2492 individuals). This is consistent with recently reported global values of *C*_mass_ (Ma et al., 2018; Tang et al., 2018). Wood C:N ratio was assigned a value of 319 g g^−1^, the global mean value of trunk C:N ratio (*n* = 544 individuals) reported by Zhang et al. (2020). Root C:N ratio was assigned the value 94 g g^−1^ (*n* = 22 sites), the median value reported by Tian et al. (2019). Although variations in wood and root C:N ratios are not negligible (Schreeg et al., 2014; Zhang et al., 2019), we treated them as constants here. Their variation appears to be more strongly controlled by phylogeny than by the environment (Zhang et al., 2020), rendering them less suitable for global upscaling with environmental covariates.

Following the finding by Deng *et al*. (2018) that NRE decreases with temperature and humidity, we fitted a linear model for NRE as a function of *T*_g_ and *D* at 184 forest sites (Deng et al., 2018; Du et al., 2020):

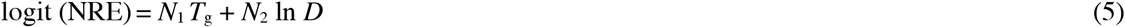

For grasslands, model selection showed that the optimal predictive model for site-mean BP (*n* = 119 sites) was fitted by *T*_g_ and gPPFD, consistent with the model fitted for forest biomes. ABP/BP (*n* = 109 sites) was non-sensitive or weakly correlated to climate predictors. Therefore, we estimated a constant value for this ratio by performing a linear regression without intercept of ABP on BP, yielding a value of 0.50. Tissue C:N ratio and NRE data for grasslands had small sample sizes, rendering fitted models insufficiently robust. Therefore, leaf and root C:N ratios were assigned constant values of 18 and 41, respectively, and NRE = 69%, all being median values across the data for grasslands.

Total N uptake (*N*_up_; g N m^−2^ yr^−1^) was estimated as:

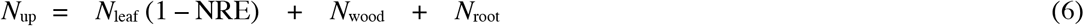

where *N*_leaf_ is BP_leaf_ divided by the leaf C:N ratio, *N*_wood_ (in forests) is BP_wood_ divided by the wood C:N ratio, and *N*_root_ is BP_root_ divided by the root C:N ratio. No resorption of N stored in roots was considered.

### Global mapping

The empirical models, developed based on site-specific observations, were applied globally – driven by global gridded data – in order to obtain up-scaled estimates of all component fluxes. Global maps were initially created for forests and grasslands separately (see Equations 3-6). The MODIS IGBP land cover map (Sulla-Menashe & Friedl, 2018) was then used to determine the fraction of forests versus grasslands in each grid cell. Six land-cover classes (evergreen needleleaf, evergreen broadleaf, deciduous needleleaf, deciduous broadleaf, mixed forests and shrublands) were treated as forest, and one as grassland. The final value assigned to each grid cell was a weighted average of the values for forests and grasslands; weighting was based on the relative cover of forests versus grasslands, normalized to a sum of 100% (thus disregarding, e.g., urban or agricultural land in order to focus on natural and semi-natural vegetation). The uncertainties of global estimations of C and N uptake were computed using standard error propagation methods. Negative BP values arising in a few cases (0.2% and 1.4% of global grid cells in forest and grassland respectively) were ignored. Global NUE was calculated based on area-weighted BP and *N*_up_ in each grid-cell.

### Factors contributing to NUE and *N*_up_

We conducted a variable importance analysis using the Lindeman, Merenda and Gold (LMG) statistic (Grömping, 2006) for *N*_up_ and NUE separately in relation to all predictor variables, based on the global gridded data. LMG statistics were calculated only for the forest models because N cycling rates in grassland depended only on *T*_g_ and gPPFD and showed little variation.

### Comparison with global vegetation models

We analysed global simulations, by 12 dynamic global vegetation models (DGVMs) in version 8 of the TRENDY model ensemble, driven by varying CO_2_ and climate but with fixed (pre-industrial) land use (the S2 simulation protocol). The same simulations were also contributed to the annual Global Carbon Budget publication for 2019 (Friedlingstein et al., 2019). The models do not distinguish BP from NPP, so we compared our BP values with DGVM-simulated NPP (variable ‘NPP’ in TRENDY outputs). We also compared our *N*_up_ values with DGVM-simulated *N*_up_ (variable ‘fN_up_’ in TRENDY outputs). These comparisons were made using values extracted from global simulations for the same sites as in other analyses.

Since all the TRENDY DGVMs and our statistical models used climate variables as predictors, we also compared partial residual relationships based on linear regressions for predicted site-level BP, *N*_up_ and NUE in relation to *T*_g_, gPPFD and *D*.

## RESULTS

### Predicting component fluxes

Component fluxes showed stronger environmental dependencies, and more accurate predictions, in forests than in grasslands.

In forests (Table 1, Fig. 1), BP and ABP/BP both decreased with increasing gPPFD (growing season mean PPFD), soil C:N ratio, and stand age; but increased with growth temperature. (Note that gPPFD tends to be greater if the growing season is shorter, thus decreasing BP.) BP also increased with fAPAR. This is expected because photosynthesis depends on canopy light absorption. The ratio BP_leaf_/ABP increased with fAPAR and gPPFD, but decreased with aridity. The regression models explained 44%, 18% and 8% of observed variance in forest BP, ABP/BP and BP_leaf_/ABP respectively (Table 1). Analyses using directly measured (instead of mapped) values of predictors showed broadly consistent patterns – with the exceptions that BP was not significantly related to measured soil C:N ratio, and ABP/BP was not significantly related to gPPFD (Fig. S5), probably due to the reduced sample sizes.

**Table 1.** Empirical models and constants. Fitted models or constants are shown for biomass production (BP; g C m^−2^ yr^−1^); the ratio of above-ground biomass production (ABP, g C m^−2^ yr^−1^) to BP; the ratio of leaf biomass production (BP_leaf_, g C m^−2^ yr^−1^) to ABP; leaf nitrogen per unit mass (*N*_mass_, unitless); leaf carbon per unit mass (*C*_mass_, unitless); root carbon-to-nitrogen ratio (C:N, g C g^−1^ N); wood carbon-to-nitrogen ratio (C:N, g C g^−1^ N) and nitrogen resorption efficiency (NRE, unitless). Site-level predictors were all mapped values (see Fig. S2): soil C:N ratio (C:N, g C g^−1^ N), forest stand age (age, years), fraction of absorbed photosynthetically active radiation (fAPAR, unitless), incident photosynthetic photon flux density averaged over the growing season (gPPFD, μmol m^−2^ s^−1^), growth temperature (*T*_g_,°C), vapour pressure deficit (*D*, kPa), maximum rate of carboxylation at 25°C (*V*_cmax25_, μmol m^−2^ s^− 1^) and leaf mass per unit area (LMA, g m^−2^). In forests, ABP/BP, BP_leaf_/ABP and NRE were logit-transformed and soil C:N ratio, forest stand age, gPPFD and *D* were log-transformed. Partial residual plots are presented in Fig. 1.

| Response variables | Fitted regressions | $R^2$ |
| --- | --- | --- |
| **Forest** |  |  |
| BP | $2837 + 15.7 T_g - 277 \ln \text{gPPFD} - 377 \ln \text{C:N} - 91.2 \ln \text{age} + 861 \text{fAPAR}$ | 0.44 |
| logit ABP/BP | $15.0 + 0.0266 T_g - 1.77 \ln \text{gPPFD} - 1.44 \ln \text{C:N} - 0.128 \ln \text{age}$ | 0.18 |
| logit $\text{BP}_{\text{leaf}}$/ABP | $-12.0 + 1.83 \ln \text{gPPFD} + 1.01 \text{fAPAR} - 0.486 \ln D$ | 0.08 |
| Leaf $N_{\text{mass}}$ | $0.00584 V_{\text{cmax}25}/\text{LMA} + 0.0161$ | 0.21 |
| Leaf $C_{\text{mass}}$ | 0.47 (median of 79 sites) | |
| Root C:N ratio | 94 (median of 22 sites) |  |
| Wood C:N ratio | 319 (mean of 544 individuals) |  |
| logit NRE | $1.15 - 0.0544 T_g + 0.282 \ln D$ | 0.23 |
| **Grassland** |  |  |
| BP | $4874 + 27.1 T_g - 797 \ln \text{gPPFD}$ | 0.23 |
| ABP/BP | 0.50 (fitted by regression with zero intercept) |  |
| Leaf C:N ratio | 18 (median of 215 sites) |  |
| Root C:N ratio | 41 (median of 71 sites) |  |
| NRE | 0.69 (median of 26 sites) |  |

**Fig. 1.**
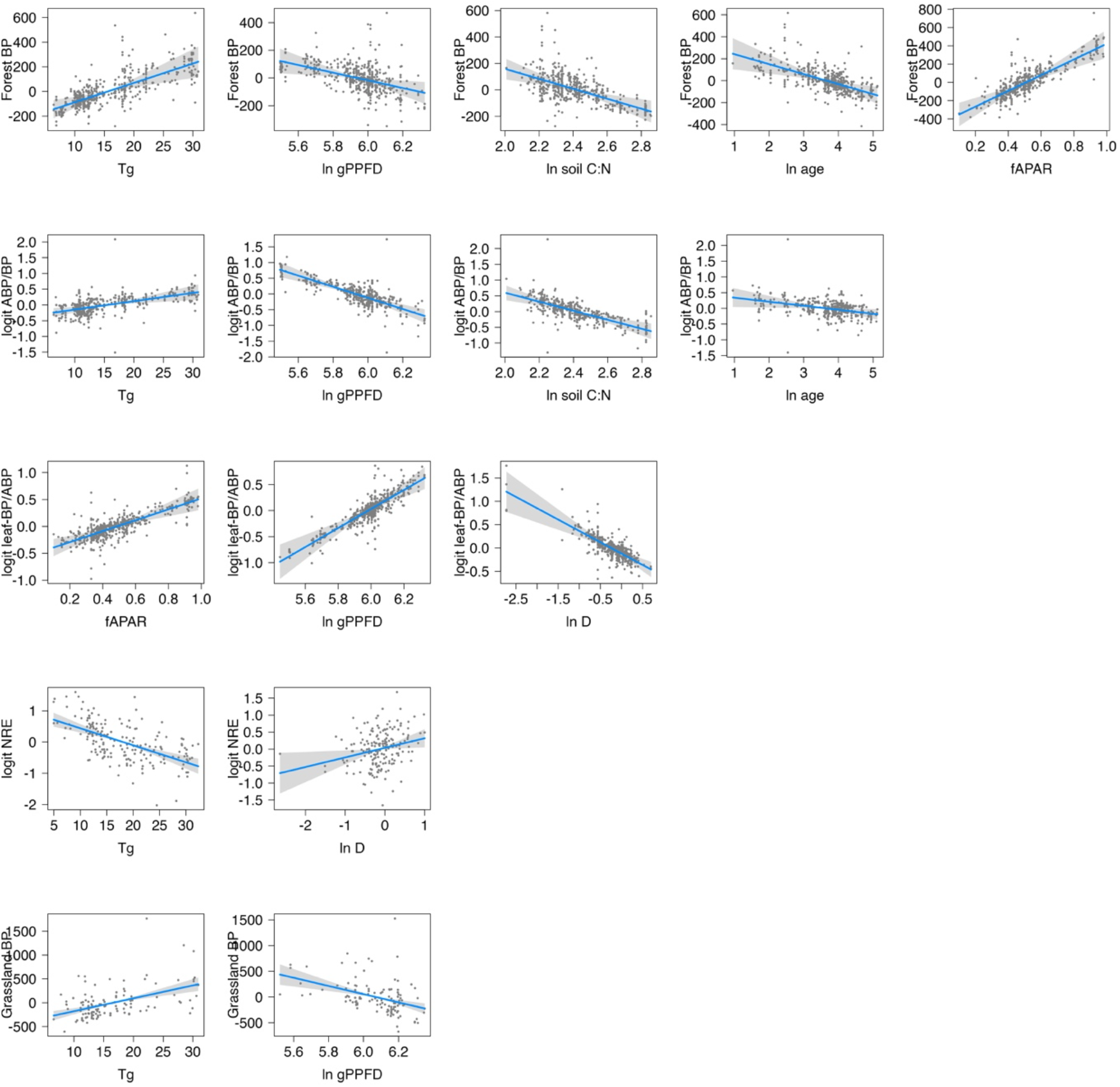
Partial residual plots for statistical models developed to predict: (1) biomass production (BP; gC m^-2^yr^-1^) in forest; (2) the ratio of aboveground biomass production (ABP; gC m^-2^yr^-1^) to BP; (3) the ratio of leaf biomass production (BP_leaf_; gC m^-2^yr^-1^) to ABP; (4) nitrogen resorption efficiency (NRE); (5) BP in grassland. Predictors are *mapped* soil C/N, stand-age, fraction of absorbed photosynthetically active radiation (fAPAR), incident photosynthetic photon flux density averaged over the growing season (gPPFD), growth temperature (*T*_g_) and vapour pressure deficit (*D*). All response variables were logit transformed, and predictors for soil C/N, age, gPPFD and *D* were log-transformed. Statistical models of forest BP, ABP/BP and BP_leaf_/ABP used linear mixed-effects models, where site is the random intercept, and each point is represented by a measured value at ecosystem level. The statistical models for NRE and grassland BP are linear regressions, with each point representing a measured site-mean value.

In grasslands, BP increased with *T*_g_ but decreased with gPPFD – qualitatively consistent with the response in forests, but explaining only 23% of observed variance. The relationship of ABP/BP to environmental variables was non-significant, hence we applied a fixed ratio ABP/BP = 0.50 (*n* = 109).

Predictions derived from the empirical models (extracted from global simulation and interpolated to site locations) showed general agreement with stand-scale measurements for BP (*R*^2^ = 0.45: Fig. 2), ABP (*R*^2^ = 0.44), BBP (*R*^2^ = 0.16), BP_leaf_ (*R*^2^ = 0.51) and BP_wood_ (*R*^2^ = 0.29) in forests; and BP (*R*^2^ = 0.24), ABP (*R*^2^ = 0.13) and BBP (*R*^2^ = 0.20) in grasslands.

**Fig. 2.**
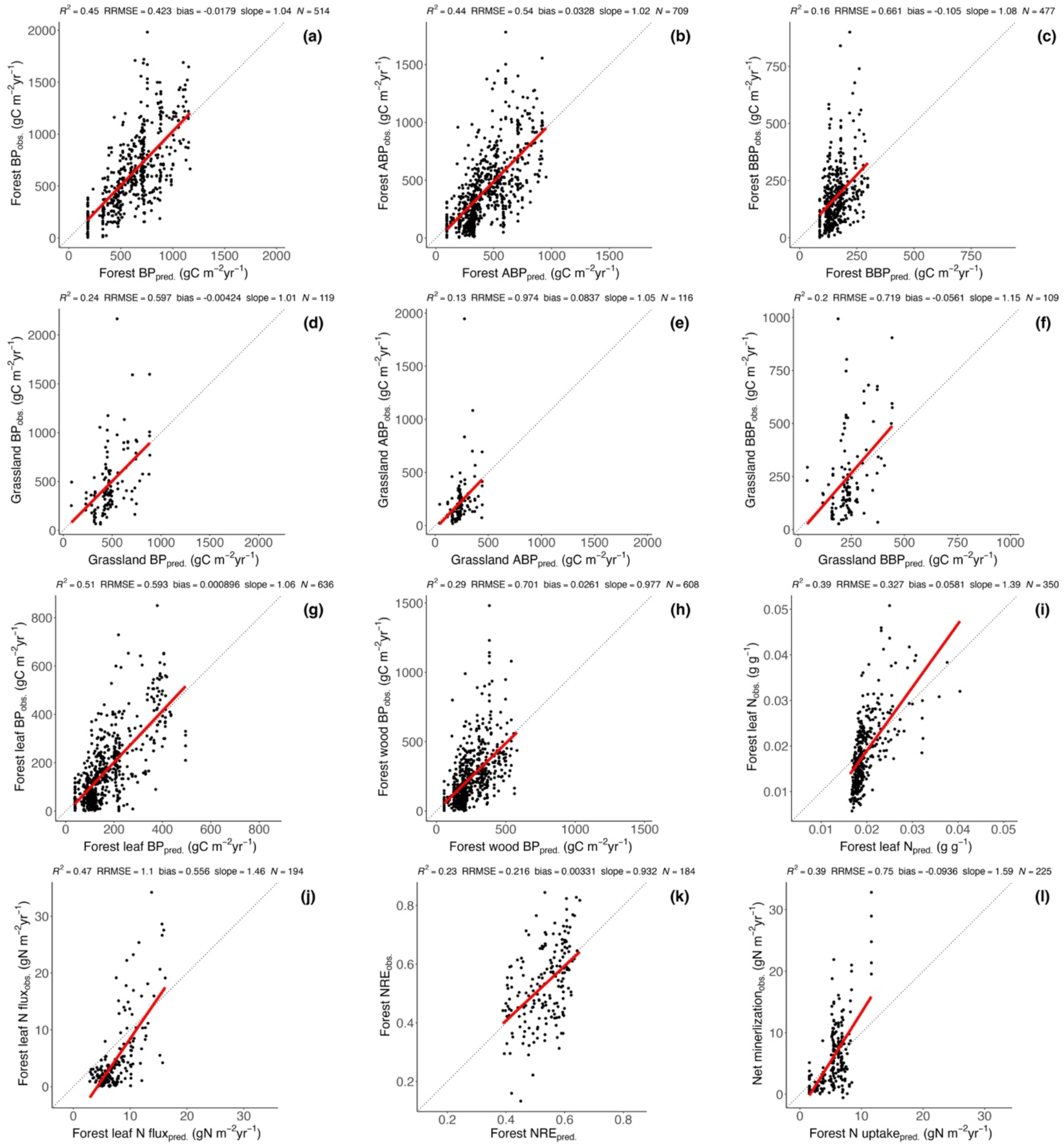
Evaluation of model predictions (see Table 1) against measurements. The dotted line is the 1:1 line. The red line represents the linear regression of modelled vs. observed values. Points in (i, k) represent site-mean values of leaf N (g g^−1^) and nitrogen resorption efficiency (NRE, unitless). Points in (a–h, j, l) represent each individual recorded for biomass production (BP, g C m^−2^ yr^−1^), above-ground biomass production (ABP, g C m^−2^yr^−1^), below-ground biomass production (BBP, g C m^−2^yr^−1^), leaf biomass production (BP_leaf_, g C m^−2^ yr^−1^), wood biomass production (BP_wood_, g C m^−2^ yr^−1^), leaf N flux (BP_leaf_ divided by leaf C:N ratio, g N m^−2^ yr^−1^) and net N mineralization (g N m^−2^ yr^−1^).

The relationship for *N*_mass_ as a linear function of *V*_cmax25_/ LMA (Eq. 5) explained 21% of the variance in observed *N*_mass_. NRE was found to decrease with higher temperature and humidity; the corresponding regression model explained 23% of observed variance (Table 1).

Finally, we estimated *N*_up_ from the combination of BP, C allocation, tissue C:N ratios and NRE, yielding good predictions of observed *N*_mass_ (*R*^2^ = 0.39, Fig. 2i) using measured *V*_cmax25_ and LMA as predictors; leaf N flux (*R*^2^ = 0.47, Fig. 2j); NRE (*R*^2^ = 0.23, Fig. 2k); and *N*_min_ (*R*^2^ = 0.39, compared to modelled *N*_up_ in Fig. 2l) in forests.

### Global carbon and nitrogen cycling

Annual global BP was estimated as 72 ± 14 Pg C yr^−1^ (Fig. 3, Table S2). Modelled BP and ABP were highest in tropical and sub-tropical forests. This is expected due to the year-round growing seasons in the tropics, and because both BP and ABP/BP are favoured by low soil C:N ratios and higher temperatures in these regions (Fig. S2). Leaf C:N ratio was also higher in tropical forests, primarily driven by low values of *V*_cmax25_ at high temperatures. NRE increased towards higher latitudes, as expected due to lower temperatures.

**Fig. 3.**
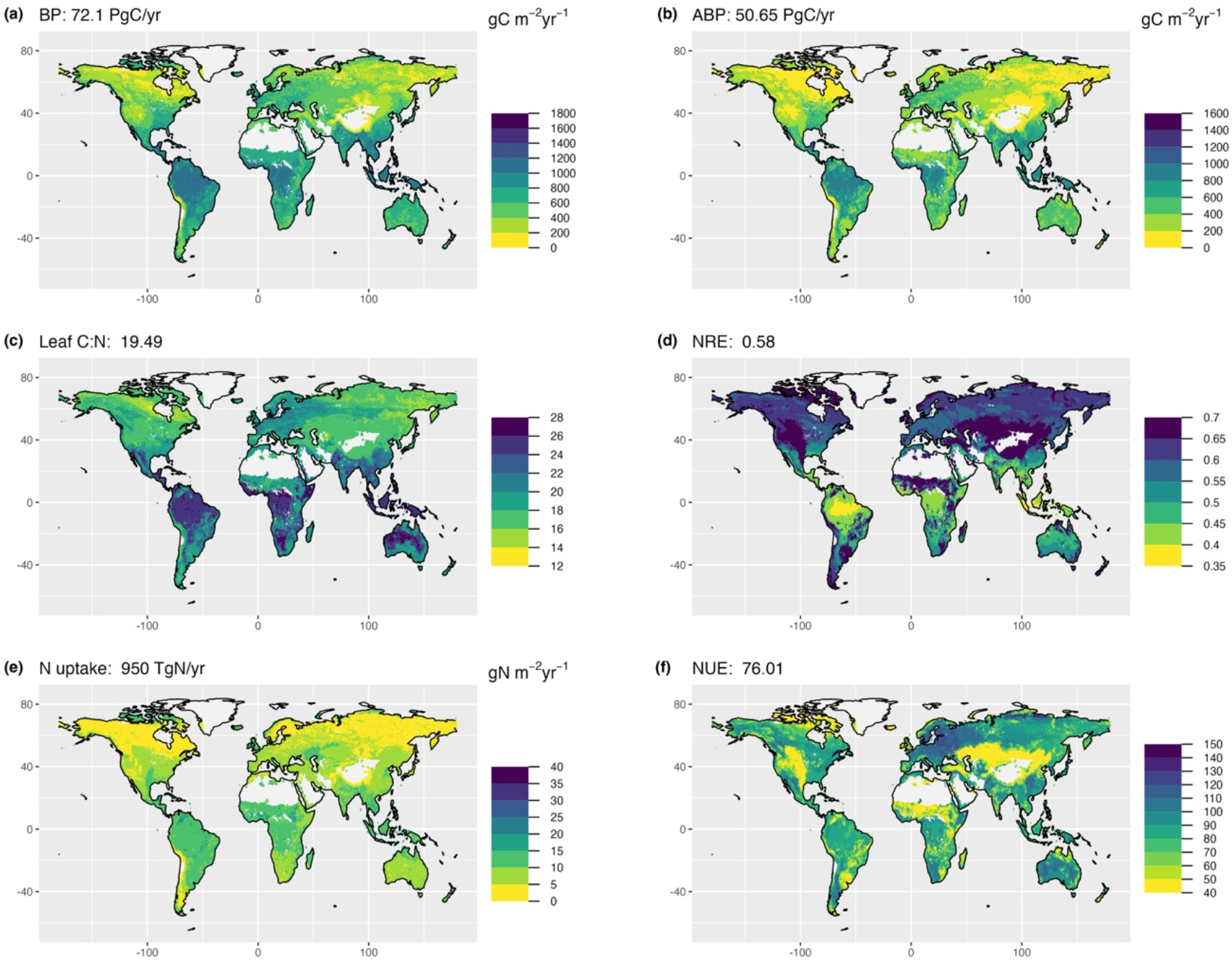
Global simulations of biomass production (BP, g C m^−2^ yr^−1^), above-ground biomass production (ABP; g C m^−2^ yr^−1^), leaf carbon-to-nitrogen ratio (leaf C:N), nitrogen resorption efficiency (NRE), N uptake (g N m^−2^ yr^−1^) and nitrogen use efficiency (NUE, the ratio of BP to N uptake). The value at the top of each panel is a global estimate. The observed sites used for fitted model and evaluation are shown in Fig. S3.

Annual global *N*_up_ was estimated as 950 ± 260 Tg N yr^−1^, with a global pattern similar to that of BP. Global NUE was estimated as 76 ± 26 g C/g N. Global mean forest NUE was estimated as 91 ± 37 g C/g N, determined by multiple climatic and soil factors. The global pattern of forest NUE differed from that of BP or *N*_up_, increasing from tropical to boreal forests. Grassland NUE was assigned a constant value of 48 ± 36 g C/g N, as we were unable to estimate the environmental dependencies of most components in grasslands (Table 1).

### Environmental dependencies of *N*_up_ and NUE

The data-driven models developed here allowed us to quantify the importance of different component processes for *N*_up_ and NUE in forests. According to the LMG statistics, variations in BP, allocation, NRE and leaf N:C ratio and NRE respectively explained 45%, 22%, 22% and 11% of variation in modelled forest *N*_up_. Climate variables (*T*_g_, *D*, gPPFD) and (independently modelled, but entirely climate-driven) *V*_cmax25_ together explained 57% of variation in modelled forest *N*_up_. fAPAR, soil C:N ratio, stand age and LMA respectively explained 28%, 10%, 5% and 0.6%.

Variations in allocation, leaf N:C, NRE and BP respectively explained 71%, 13%, 11% and 5% of the modelled variation in NUE. Climate variables and climate-derived *V*_cmax25_ together explained 76% of the modelled variance in NUE. fAPAR, LMA, stand-age and soil C:N respectively explained 16%, 5%, 2% and 1%.

Overall, we found NUE in forests to increase with LMA, age, soil C:N ratio and aridity; and to decrease with fAPAR, *T*_g_, *V*_cmax25_ and gPPFD (Table 1). The pattern of variation in NUE is dominated by climate via its effects on biomass allocation – especially allocation to leaves, which are richer in N than other tissues. Increasing leaf allocation is the primary factor leading to decreasing NUE (Fig. S6). Thus, NUE decreases with temperature because lower temperatures decrease aboveground allocation, including allocation to leaves; and it increases with aridity because leaf allocation is reduced in dry climates. These two patterns are compounded by the effects of NRE, which is greater in both drier and colder environments, leading to increased NUE. The decrease of NUE with gPPFD is also primarily driven by leaf allocation: increasing gPPFD decreases the ratio ABP/BP, but more importantly, increases the ratio BP_leaf_/ABP, thereby reducing NUE.

Aboveground allocation was also reduced in soils with higher C:N ratios. However, soil C:N ratio accounted for only 1% of modelled variance in NUE, an effect much smaller than that of climate.

NUE in grassland was assigned a globally fixed value, but this value is lower than that of forests due to the high C:N ratio of wood. Low NUE in grassland explains the relatively sharp transitions (seen in Fig. 3f) between low values in semi-arid grasslands and much higher values in nearby dry forests.

### Comparison with global vegetation models

Comparing our global estimates with measurements for BP in forest yielded *R*^2^ = 0.45 (Fig. 2a). Comparison of our global estimates of *N*_up_ with *N*_min_ data yielded *R*^2^ = 0.39 (Fig. 2l). TRENDY models performed variably in comparison with these measurements (Table 2). Many models showed good performance, approaching that of our benchmark model, for BP. Among the four models allowing comparison with N uptake, however, none shows *R*^2^ greater than half that of our benchmark.

**Table 2.** Statistics for the comparison of global simulations (from our study and TRENDY output) interpolated to measurement sites. Measured variables are biomass production (BP) and net mineralization (N_min_). *R*^2^ is the coefficient of determination; RRMSE is the relative root-mean-square error, as a proportion of the observed mean value; ‘Rel. bias’ is the difference between observed and predicted mean values, expressed as a proportion of the observed mean value; Slope is the slope of the linear regression of observed against predicted values. The site distribution is shown in Fig. S3.

| **Predicted BP vs. measured BP** |  |  |  |  |
| --- | --- | --- | --- | --- |
| | $R^2$ | RRMSE | Rel. bias | Slope |
| **Our model** | **0.45** | **0.42** | **−0.02** | **1.04** |
| CABLE (Haverd et al., 2018) | 0.37 | 0.51 | 0.10 | 0.64 |
| ISAM (Meiyappan et al., 2015) | 0.36 | 0.52 | −0.25 | 0.98 |
| ISBA (Decharme et al., 2019) | 0.30 | 0.49 | −0.10 | 0.84 |
| JULES (Sellar et al., 2019) | 0.27 | 0.61 | 0.26 | 0.56 |
| LPJ (Smith et al., 2014) | 0.05 | 0.63 | −0.23 | 0.45 |
| ORCHIDEE (Goll et al., 2017) | 0.42 | 0.49 | −0.21 | 0.92 |
| ORCHICNP (Krinner et al., 2005) | 0.12 | 0.57 | −0.02 | 0.50 |
| SDGVM (Walker et al., 2017) | 0.19 | 0.53 | −0.12 | 0.98 |
| CLASS (Melton & Arora, 2016) | 0.22 | 0.67 | 0.32 | 0.48 |
| CLM (Lawrence et al., 2019) | 0.19 | 0.53 | −0.01 | 0.70 |
| JSBACH (Mauritsen et al., 2019) | 0.31 | 0.71 | 0.23 | 0.41 |
| LPX (Lienert & Joos, 2018) | 0.32 | 0.50 | −0.12 | 0.74 |
| **Predicted N uptake vs. measured net mineralization** |  |  |  |  |
| **Our model** | **0.39** | **0.75** | **−0.09** | **1.59** |
| ISAM (Meiyappan et al., 2015) | 0.08 | 0.92 | −0.20 | 0.54 |
| ORCHICNP (Krinner et al., 2005) | 0.13 | 1.19 | 0.53 | 0.56 |
| JSBACH (Mauritsen et al., 2019) | 0.09 | 1.21 | 0.34 | 0.24 |
| LPX (Lienert & Joos, 2018) | 0.14 | 0.97 | 0.45 | 0.74 |

We also compared the climatic dependencies of our global estimates of BP, *N*_up_ and NUE with TRENDY DGVMs. All the DGVMs captured the positive response of BP to temperature. Most also captured the decrease of BP with gPPFD (Fig. 4). The representation of global patterns for *N*_up_ and NUE in relation to climate, however, showed a diversity of responses. One model showed the wrong sign for the temperature dependency of both *N*_up_ and NUE.

**Fig. 4.**
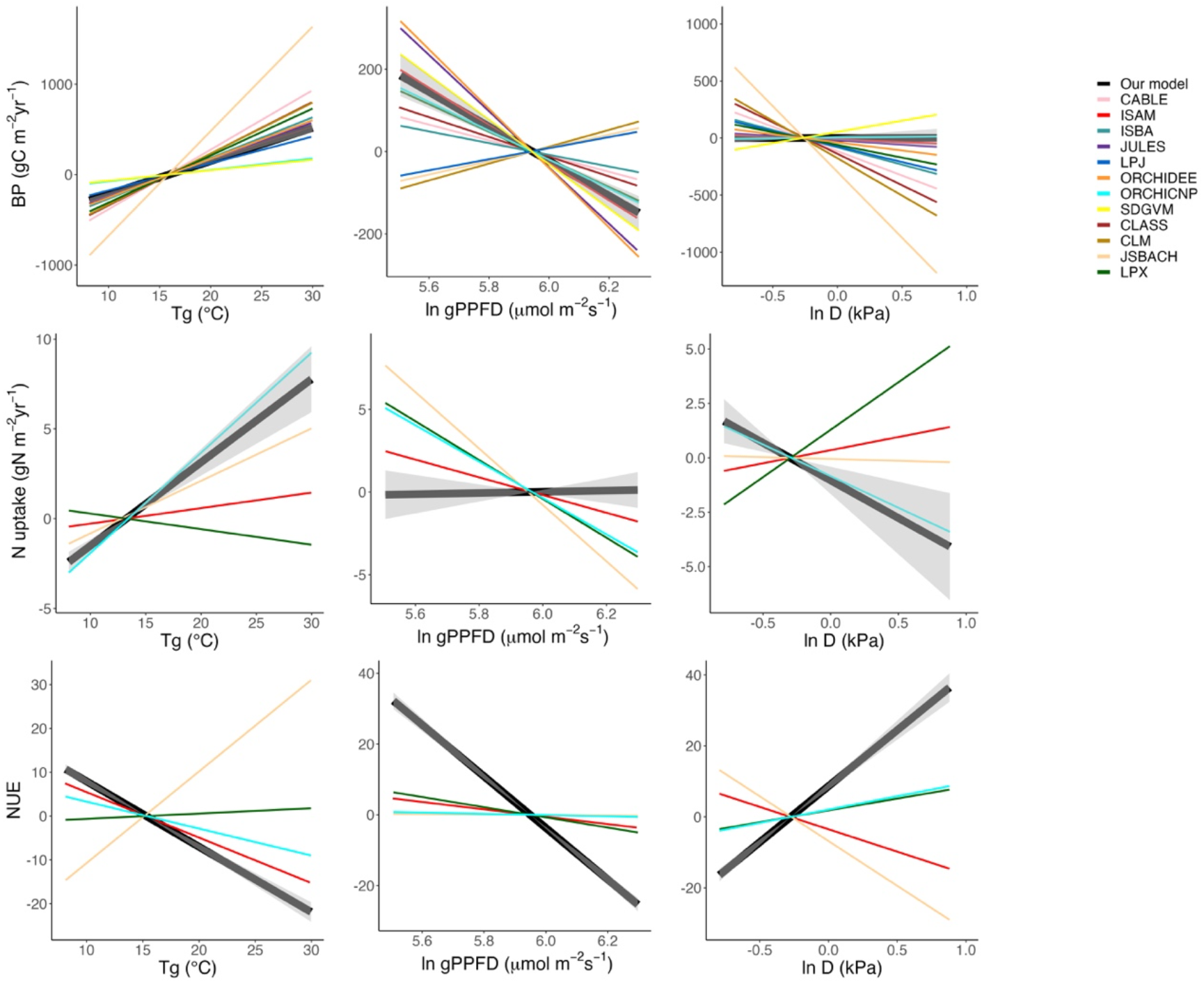
Partial residual relationships for modelled biomass production (BP, g C m^−2^ yr^−1^), N uptake (*N*_up_, g N m^−2^ yr^−1^) and nitrogen use efficiency (NUE, g C g^−1^ N) as functions of incident photosynthetic photon flux density averaged over the growing season (gPPFD, μmol m^−2^ s^−1^), growth temperature (*T*_g_,°C) and vapour pressure deficit (*D*, kPa) during the thermal growing season. Predicted site-level BP was based on all measured BP plots in forests; predicted site-level *N*_up_ was based on all measured N mineralization plots in forests; predicted site-level NUE was based on all measured BP and N mineralization plots in forest (Fig. S3a, e).

## DISCUSSION

One recent study suggested that all forest C fluxes (autotrophic respiration, NPP, above-and below-ground NPP) display similar trends with respect to latitude, temperature and growing-season length (Banbury Morgan et al., 2021), with no difference in allocation at the global scale. Many observational and experimental studies contradict this, indicating that C allocation is influenced by climate and soil factors including light (Poorter et al., 2012), water (Ma et al., 2021; Schenk & Jackson, 2002; Zhang et al., 2019), temperature (Lambers et al., 2008; Ma et al., 2021), CO_2_ (Poorter et al., 2021; Terrer et al., 2018) and nutrient availability (Litton et al., 2007; Ven et al., 2019, 2020; Yan et al., 2019). Here, we focus on observed variations in BP, *N*_up_ and their ratio, NUE. Our analysis documents the differentiated responses of these three quantities to biotic and environmental factors, and the particular importance of variations in C allocation in determining NUE.

BP is shown to be positively related to growth temperature and light absorption, while declining with the seasonal concentration of light availability – features captured by most of the DGVMs. Additional controls on BP are soil C:N ratio (with more organic soils supporting lower BP), and forest stand age. N supply limitation on BP is well supported by observational studies (LeBauer & Treseder, 2008; Vicca et al., 2012). BP increases towards lower soil C:N ratio since higher N availability increases whole-plant photosynthesis and growth (Vicca et al., 2012). Soil C:N ratio is a relatively crude proxy for N availability (Maire et al., 2015) but it emerged here as a significant control on BP, in line with previous research (Radujković et al., 2021; Terrer et al., 2019; Van Sundert et al., 2020; Vicca et al., 2018). Regarding stand age, the longer transport pathway for water in taller trees can result in reduced stomatal conductance and photosynthesis (Drake et al., 2011) while the greater sapwood mass is required to support a given leaf area and implies increased maintenance respiration (Collalti & Prentice, 2019; Mori et al., 2010; Reich et al., 2008) – both effects potentially contributing to a decline in BP.

BP itself emerged as the most important predictor of *N*_up_ in our analysis – inevitably, given that *N*_up_ has to match the stoichiometric requirements of plant growth (Cleveland et al., 2013; Zaehle et al., 2014). This principle is therefore built into our calculation of *N*_up_. Given the strong climatic controls of BP, it also follows that climate exerts a primary control on *N*_up_. The involvement of soil C:N ratio as a secondary control of modelled *N*_up_ is consistent with soil N limitations on whole-plant C and N uptake (Lawrence et al., 2019; Mauritsen et al., 2019).

The ratio of ABP to BP showed responses to the environment that are qualitatively similar to those of BP, including similar responses to soil C:N ratio – indicating that less fertile soil conditions tend to increase BBP relative to ABP. From high to low soil N availability, as indicated here by increasing soil C:N ratio, increasing allocation of C to roots is commonly observed along with decreasing allocation to above-ground production (Franklin et al., 2012; Peng et al., 2017). The large below-ground C allocation in soils with low N availability helps ectomycorrhizal (ECM) fungi to acquire N from soil organic matter (Phillips et al., 2013), causing accelerated soil C turnover (Pregitzer et al., 2008) and N cycling (Zak et al., 2011).

The ratio of BP_leaf_ to ABP in forests increases with moisture, here measured by the growing season mean vapour pressure deficit, probably because sapwood area per unit leaf area increases with aridity due to the additional water requirement of a given rate of photosynthesis under dry conditions (Mencuccini & Grace, 1995). The positive effect of fAPAR on this ratio is expected, due to the direct link between leaf light absorption and photosynthesis. The positive effect of gPPFD may reflect the fact that the annual C allocation to leaves is determined by the annual maximum foliage: a shorter growing season allows each unit of leaf carbon to produce less photosynthate, and therefore likely less BP.

Tissue stoichiometry and NRE explained less variance in *N*_up_ than BP and biomass allocation (Wang et al., 2018b). Modelled leaf C:N ratios decreased towards high latitudes, driven by *V*_cmax25_ increasing towards cold climates (Peng et al., 2021). This pattern is consistent with the principle of the LPJ-GUESS model (Smith et al., 2014), which assumes that leaf N is driven by its climate-driven demand rather than soil N supply. It is not captured by models wherein leaf C:N ratio is assigned PFT-specific (Oleson *et al*., 2010; Lawrence *et al*., 2019) or globally fixed (Wiltshire et al., 2021) values.

Global mean NRE was shown to increase from low to high latitudes, driven by negative effects of temperature and humidity on NRE. Since rates of N cycling are enhanced in warmer and wetter environments, N supply from resorption is relatively less important in tropical regions relative to higher latitudes (Deng et al., 2018; Du et al., 2020).

Our analysis casts some light on the opposition between ‘biogeochemical niche differentiation’ (Peñuelas et al., 2019; Sardans et al., 2021) and ‘climate-driven demand’ (Wang et al., 2018b) as the primary controls of ecosystem stoichiometry. Globally, soil C:N ratio accounted for just 10% of the modelled variation in *N*_up_, and 1% of the modelled variation in NUE. N deposition was initially considered as an additional predictor for BP and allocation, but produced no improvement in the fitted model performance for *N*_up_ and was therefore discarded for the further analysis and modelling. By contrast, climate accounted for 57% of modelled variation in *N*_up_, and 76% of modelled variation in NUE. These results point to a dominant control of *N*_up_ and NUE by climate, with a secondary influence by soils.

Overall, biomass allocation emerged as the dominant process controlling of forest NUE (driving 71% of modelled variance). This finding is consistent with GOLUM-CNP (Wang et al., 2018b), where biomass allocation explained >80% of modelled variation in NUE (defined as the ratio of gross primary production to *N*_up_ in their study). According to our models, forest NUE is primarily driven by climate, increasing towards colder and drier climates. A possible interpretation of this pattern invokes the concept that high efficiency of resource use is favoured when resource supplies are more limiting to production (Harrington et al., 2001). This interpretation is supported by observations (Gill & Finzi, 2016) and simulations (Wang et al., 2018b), indicating that NUE increases from tropical to boreal forest. In boreal forests, much of the N pool is bound to organic material and is depolymerized by microbially produced hydrolytic and oxidative enzymes whose activity is limited by low temperatures (Gill & Finzi, 2016). To overcome this limitation, boreal plants are dependent on ECM or ericoid microbial symbioses for efficient N acquisition (Högberg et al., 2010; Näsholm, 1998; Terrer et al., 2019), requiring greater C allocation below ground (Gill & Finzi, 2016). The increase of NUE with aridity can be explained by reduced allocation to (N-rich) leaves. NRE also plays a role, as N conservation (by resorption) is a favoured strategy in colder and drier environments.

Globally, however, the lowest NUE according to our mapping is encountered in arid regions (including Central Asia, the interior West of North America and the Sahel) where grasslands dominate. The highest NUE is shown primarily in temperate forests, especially in northern Europe and China, where relatively low temperatures increase belowground allocation and decrease NRE. N deposition in these regions is among the highest globally (Reay et al., 2008), suggesting that N deposition is not a primary control on NUE. Much of Australia is also shown as a region of high NUE due to the occurrence of forests in dry climates that favour reduced leaf allocation and high NRE.

The increase of NUE from tropical to boreal forest has also been linked to decreasing soil N supply or soil N:P ratio (Gill & Finzi, 2016). In our analysis soil C:N ratio accounted for 10% of explained variance for *N*_up_ but only 1% for NUE, suggesting that soil N supply is a minor factor determining NUE and implying that effects of climate – which include indirect effects, such as the increased C cost of N acquisition at low temperatures – are dominant.

Existing coupled C-N cycle models predict plant C allocation by a variety of methods. Some assume fixed (PFT-specific) allocation fractions; others embed functional relationships between different dimensions, such as leaf and sapwood area (Zaehle et al., 2014), in more process-based formulations. Some analyses have used satellite observations of LAI and aboveground biomass (CARDAMOM; Bloom et al., 2016) directly as inputs. Given the importance of C allocation for the N cycle, it is important to check that assumptions made about C allocation are realistic. This is not always the case. For example, Wang et al. (2018) noted that in CARDAMOM, allocation to wood production was commonly >60% (this is rare in measurements) and that leaf turnover time of leaves in temperate and boreal biomes was < 1yr (but for evergreen leaves it is commonly 2.5 to 10yr). Different assumptions about C allocation will necessarily lead to divergent estimations of NUE, so the present situation implies huge uncertainty about the patterns and controls of NUE that could be reduced by systematic comparison against observationally based benchmarks.

We have noted that the distinction of forest and grassland distributions is an important factor determining NUE in our global maps. Our approach is consistent with TRENDY models that assign PFT-specific leaf C:N ratios, C allocation fractions and NRE (Smith et al., 2014; Wiltshire et al., 2021). Nonetheless, it is simplistic in (a) assuming sharp boundaries and (b) assigning fixed values to several parameters for grasslands. More work is required to improve the estimation of plant properties related to N cycling in non-forest biomes.

Global BP was estimated as 72 ± 14 Pg C yr^-1^, larger than in earlier studies by Cleveland *et al*. (2013) (44.35 Pg C yr^−1^) and Wang *et al*. (2018) (52.50 Pg C yr^−1^). Global total *N*_up_ was estimated as 950 ± 260 Tg N yr^−1^. This falls towards the upper end of the range of estimates by six models of global C and N cycling: 465 (Wiltshire et al., 2021), 728 (Smith et al., 2014), 831 (Winkler, et al., 2017), 968 (Oleson *et al*., 2010), 1172 Tg N yr^−1^ (Lawrence et al., 2019) and 1197 Tg N yr^−1^ (Cleveland et al., 2013). Global NUE was estimated as 76 ± 26 g C g^−1^ N. This falls within the range of 50 g C g^−1^ N estimated by the TEM model (Melillo et al., 1993), 52 g C g^-1^ N by the O-CN model (Zaehle et al., 2010), 56 g C g^−1^ N by the ORCHICNP model (Goll et al., 2017) and 80 g C g^−1^ N by the ISAM model (Meiyappan et al., 2015). Thus, our central estimates of global *N*_up_ and NUE are within the broad ranges simulated by current models.

In conclusion, our data-driven modelling approach has generated quantitative relationships that are broadly consistent with experimental and observational evidence for the controls of different processes contributing to the coupling of the terrestrial C and N cycles. Data on non-forest ecosystems are however sparse, limiting the information that can be derived from them. Further limitations of this study include relatively low *R*^2^ values for some comparisons (especially those related to C allocation to different tissues), and the lack of available measurements of soil factors more directly related to plant function than soil C:N ratio, a relatively crude metric of N availability. Despite these limitations, our analysis provides new benchmarks for coupled C–N cycle modelling. We have presented initial comparisons with TRENDY model simulations, based on their publicly available outputs. Simplified representations of allocation and tissue C:N ratios in TRENDY models may be responsible for divergence of the modelled *N*_up_ and NUE responses to climate from one another, and from the benchmarks provided here. A more in-depth examination of model behaviour would be worthwhile in light of our findings, potentially contributing to a reduction in the uncertainties associated with N cycle constraints on ecosystem C uptake in a changing environment.

## Acknowledgements

We acknowledge TRENDY v8 modellers, namely the late V Haverd (CABLE), A Jain (ISAM), E Joetzjer (ISBA), A Wiltshire (JULES), B Poulter (LPJ), V Bastrikov (ORCHIDEE), P McGuire (SDGVM), V Arora (CLASS), D Lombardozzi (CLM), J Nabel (JSBACH) and S Lienert (LPX) for contributing model outputs. We thank S Sitch and P Friedlingstein for organizing TRENDY. YP and BDS were funded by the Swiss National Science Foundation (grant No. PCEFP2_181115). ICP acknowledges funding from the European Research Council (ERC) under the European Union’s Horizon 2020 research and innovation programme (grant agreement No: 787203 REALM). DT was supported by National Natural Science Foundation of China (grant No: 31800397) and Young Elite Scientists Sponsorship Program by China Association for Science and Technology (2021-2023, No. 2021QNRC001). XPW acknowledges funding from the National Natural Science Foundation of China (grant No: 31870430). This work is a contribution to the LEMONTREE (Land Ecosystem Models based On New Theory, obseRvations and ExperimEnts) project, funded through the generosity of Eric and Wendy Schmidt by recommendation of the Schmidt Futures program. ICP and BDS acknowledge support from this project.

## Author contributions

BDS and ICP proposed the topic and supervised the research. YP carried out all analyses, created the graphics and wrote the first draft of the manuscript. All authors provided data for the analysis and contributed to the interpretation of results and revisions of the manuscript.

## Data availability

No new data were used in the analysis and models presented here. The observed data for biomass production, C and N allocation, tissue C:N ratios and N resorption efficiency were collected from the cited publications. All measured and predicted data used in this manuscript will be publicly available once published. Code used for analyses is available via Zenodo: Yunke Peng, & Benjamin Stocker. (2022). yunkepeng/CNuptake_MS: CNuptake MS for submitted version (3.0). Zenodo. https://doi.org/10.5281/zenodo.7271823

