## Supporting information for "Global terrestrial nitrogen uptake and nitrogen use efficiency"

Table S1. Model selection for stepwise forward regression.

| Response variables | Selected predictors (* if significant) | $R^2$ | AIC | BIC |
|---|---|---|---|---|
| **BP in forest** | $T_g$* | 0.31 | 7143 | 7160 |
| | $T_g$* + fAPAR* | 0.35 | 7106 | 7127 |
| | $T_g$* + fAPAR* + ln age* | 0.41 | 7064 | 7089 |
| | $T_g$* + fAPAR* + ln age* + ln C/N* | 0.43 | 7043 | 7073 |
| | **$T_g$* + fAPAR* + ln age* + ln C/N* + gPPFD*** | **0.44** | 7027 | 7061 |
| | $T_g$* + fAPAR* + ln age* + ln C/N* + gPPFD + ln $D$ | 0.44 | **7017** | **7055** |
| **logit ABP/BP** | ln C/N* | 0.08 | 1047 | 1064 |
| **in forest** | ln C/N* + ln gPPFD* | 0.14 | **1023** | **1044** |
| | ln C/N* + ln gPPFD* + $T_g$* | 0.16 | 1025 | 1050 |
| | **ln C/N* + ln gPPFD* + $T_g$* + ln age*** | **0.18** | 1026 | 1055 |
| | ln C/N* + ln gPPFD* + $T_g$* + ln age* + ln $D$ | 0.18 | 1027 | 1061 |
| | ln C/N* + ln gPPFD* + $T_g$* + ln age + ln $D$ + fAPAR | 0.18 | 1025 | 1063 |
| **logit $BP_{leaf}$/ABP** | ln age* | 0.02 | 1376 | 1394 |
| **in forest** | ln age* + ln gPPFD* | 0.04 | 1369 | 1391 |
| | ln age* + ln gPPFD* + ln $D$* | 0.06 | 1360 | 1386 |
| | ln age + ln gPPFD* + ln $D$* + $T_g$* | 0.08 | 1354 | 1384 |
| | ln age + ln gPPFD* + ln $D$* + $T_g$ + fAPAR | 0.08 | 1355 | 1390 |
| | ln age + ln gPPFD* + ln $D$* + $T_g$ + fAPAR + ln C/N | 0.08 | 1357 | 1397 |
| | **Second iteration (age removed)** | | | |
| | fAPAR* | 0.02 | 1373 | 1391 |
| | fAPAR* + ln gPPFD* | 0.04 | 1365 | 1387 |
| | **fAPAR* + ln gPPFD* + ln $D$*** | **0.08** | **1343** | **1369** |
| | fAPAR* + ln gPPFD* + ln $D$* + ln C/N | 0.08 | 1346 | 1377 |
| | fAPAR + ln gPPFD* + ln $D$* + ln C/N + $T_g$ | 0.08 | 1353 | 1389 |
| **BP** | $T_g$* | 0.10 | 1699 | 1707 |
| **in grassland** | $T_g$* + ln gPPFD* | 0.23 | **1682** | **1694** |
| | $T_g$* + ln gPPFD + ln $D$ | 0.24 | 1683 | 1697 |
| | $T_g$* + ln gPPFD + ln $D$ + fAPAR | **0.25** | 1683 | 1700 |
| | $T_g$* + ln gPPFD + ln $D$ + fAPAR + ln C/N | 0.24 | 1685 | 1705 |

Table S2. Simulated global C and N uptakes. Units are PgC yr$^{-1}$ or PgN yr$^{-1}$.

| | **Forest** | **Grassland** | **Total** | **Literature models** |
|---|---|---|---|---|
| **BP** | 55.74 ± 10.62 | 16.36 ± 9.72 | 72.10 ± 14.40 | 44.35 (Cleveland *et al*. 2013)<br>52.50 (Wang et al. 2018)<br>56.82 (LPJ)<br>66.50 (SDGVM) |
| **ABP** | 42.47 ± 8.68 | 8.18 ± 4.86 | 50.65 ± 9.95 | |
| **BBP** | 13.27 ± 13.74 | 8.18 ± 4.86 | 21.45 ± 14.57 | |
| **Leaf BP** | 16.67 ± 4.38 | | | |
| **Wood BP** | 25.80 ± 6.08 | | | |
| **Leaf N flux * (1 – NRE)** | 0.39 ± 0.16 | 0.14 ± 0.08 | 0.53 ± 0.18 | |
| **Root N flux** | 0.14 ± 0.15 | 0.20 ± 0.12 | 0.34 ± 0.19 | |
| **Wood N flux** | 0.08 ± 0.02 | | | |
| **N uptake** | 0.61 ± 0.22 | 0.34 ± 0.15 | 0.95 ± 0.26 | 0.97 (CLM4.5)<br>1.17 (CLM5)<br>0.83 (JSBACH)<br>0.47 (JULES-ES)<br>0.73 (LPJ-GUESS) |
| **NUE (gC/gN)** | 91.38 ± 37.27 | 48.11 ± 35.61 | 76.01 ± 25.71 | 50 (TEM)<br>52 (O-CN)<br>61 (ORCHICNP)<br>80 (ISAM) |

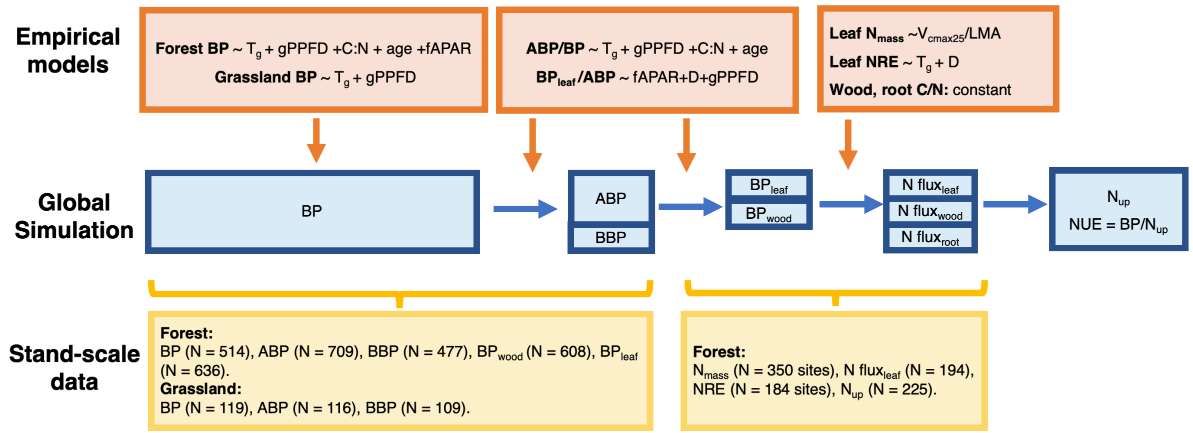


Fig. S1 Flowchart for data-driven model estimation and global simulation of carbon and nitrogen cycling. In forests and grasslands, empirical models and constants were designed to determine biomass production (BP; g C $m^{-2}$ $yr^{-1}$); the ratio of aboveground biomass production (ABP; g C $m^{-2}$ $yr^{-1}$) to BP; the ratio of leaf biomass production ($BP_{leaf}$; g C $m^{-2}yr^{-1}$) to ABP; leaf nitrogen per mass ($N_{mass}$; unitless); wood and root carbon-to-nitrogen ratio (C:N; g C/g N) and nitrogen resorption efficiency (NRE; unitless). Predictors were interpolated from global maps of soil C:N ratio (C:N; g C/g N), stand age (age; years), fraction of absorbed photosynthetically active radiation (fAPAR; unitless), incident photosynthetic photon flux density averaged over the growing season (gPPFD; µmol $m^{-2}$ $s^{-1}$), growth temperature ($T_g$; °C), vapour pressure deficit ($D$; kPa), maximum rate of carboxylation ($V_{cmax25}$, µmol $m^{-2}$ $s^{-1}$) and leaf mass per unit area (LMA, g $m^{-2}$). Global maps at 0.5° resolution were derived using data layers for soil C:N ratio (Batjes, 2015), forest stand age (Poulter *et al*., 2018), fAPAR (Pinzon & Tucker, 2014), gPPFD (Weedon *et al*., 2014), $T_g$ (Harris *et al*., 2014), $D$ (Harris *et al*., 2014), $V_{cmax25}$ (Stocker *et al*., 2020) and LMA (Moreno-Martínez *et al*., 2018) (see Fig. S2). Global simulations were tested by interpolating gridded predictions of BP, ABP, BBP, $BP_{leaf}$, $BP_{wood}$, $N_{mass}$, N $flux_{leaf}$ (defined as the product of $BP_{leaf}$ and leaf N:C ratio) and $N_{up}$ to site level.

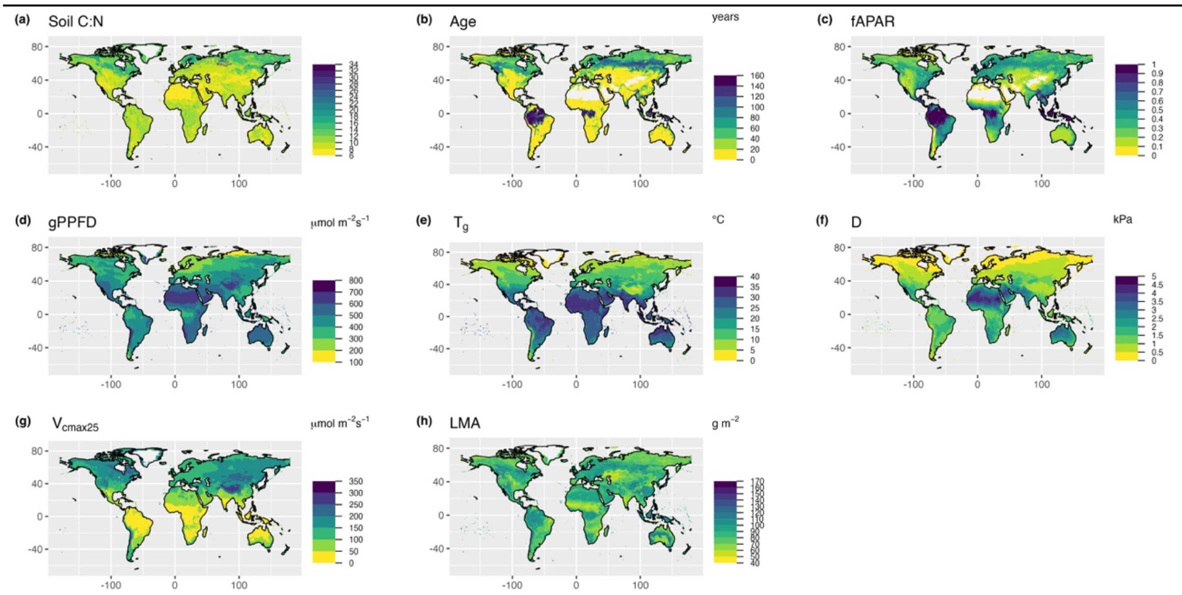


Fig.S2 All prediction fields mapped for site and global simulations: soil carbon to nitrogen ratio (C/N) (Batjes, 2015), stand-age (age) (Poulter *et al*., 2018), fraction of absorbed photosynthetically active radiation (fAPAR) (Pinzon & Tucker, 2014), incident photosynthetic photon flux density averaged over the growing season (gPPFD) (Weedon *et al*., 2014), growth temperature ($T_g$) (Harris *et al*., 2014), vapour pressure deficit ($D$) (Harris *et al*., 2014), maximum rate of carboxylation at 25˚C ($V_{cmax25}$) (Stocker *et al*., 2020) and leaf mass-per-area (LMA) (Moreno-Martínez et al. 2018).

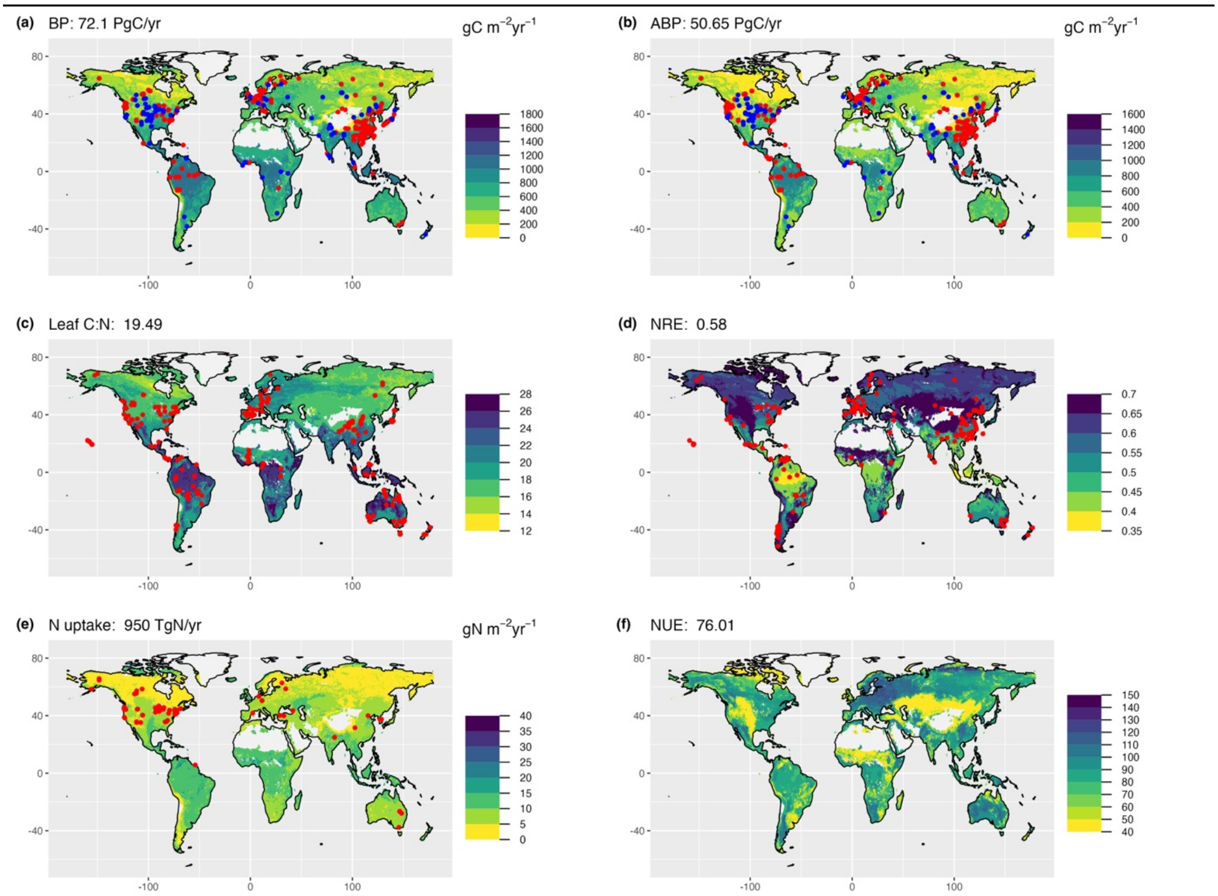


Fig. S3 Global simulations of biomass production (BP; g C m$^{-2}$ yr$^{-1}$), above-ground biomass production (ABP; g C m$^{-2}$ yr$^{-1}$), leaf carbon-to-nitrogen ratio (leaf C/ N), nitrogen resorption efficiency (NRE), terrestrial N uptake (g N m$^{-2}$ yr$^{-1}$) and nitrogen-use-efficiency (NUE). The value at the top of each panel is a global estimate. The red and blue points represent measurement sites (forests, grasslands) used for evaluation.

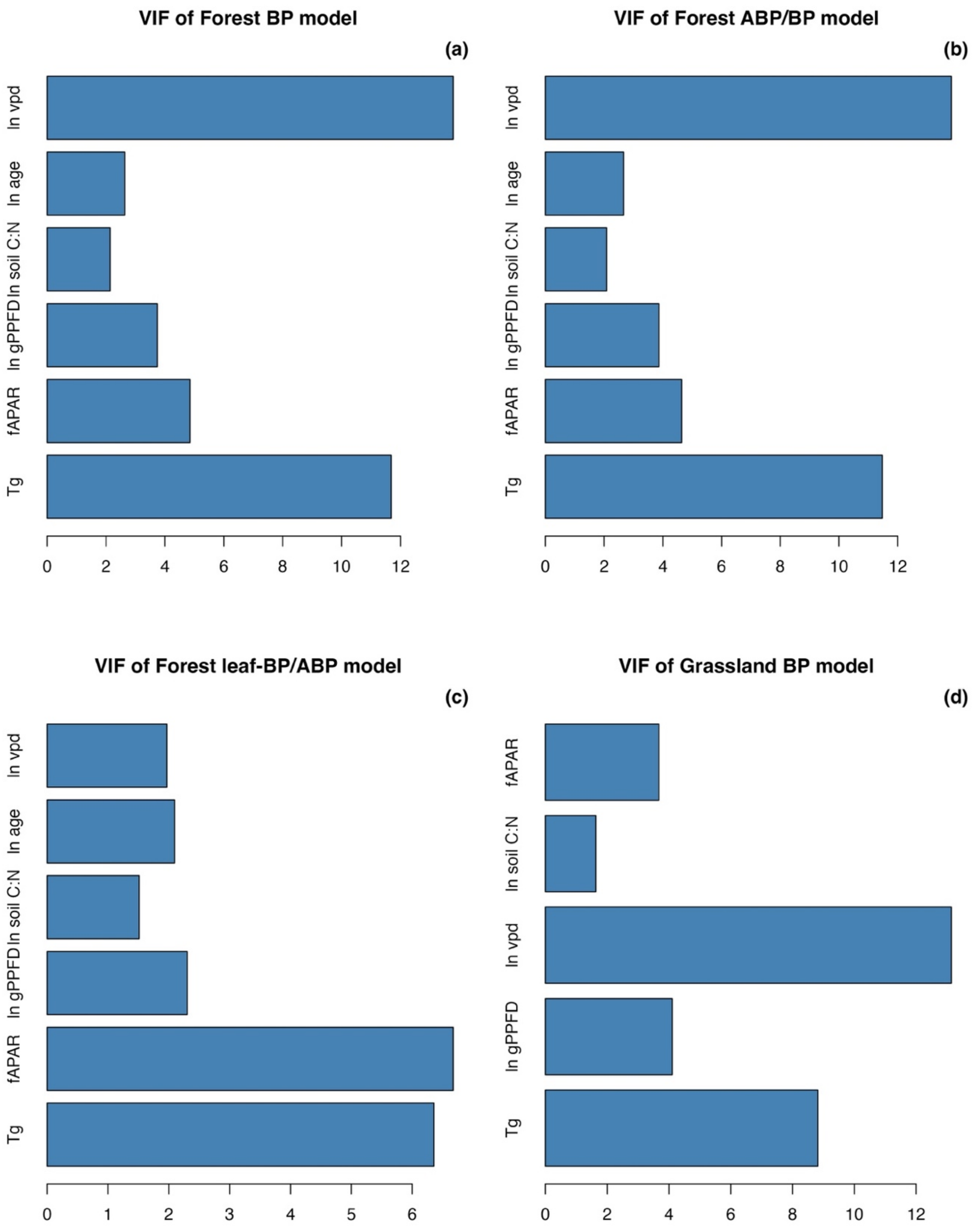


Fig. S4 Multicollinearity VIF analysis for fitted BP, ABP/BP, and leaf-BP/ABP models. Predictors are soil C/N ratio (C/N; g C/g N), forest stand age (age; years), fraction of absorbed photosynthetically active radiation (fAPAR; unitless), incident photosynthetic photon flux density averaged over the growing season (gPPFD; μmol $m^{-2}$ $s^{-1}$), growth temperature ($T_g$; °C) and vapour pressure deficit ($D$; kPa).

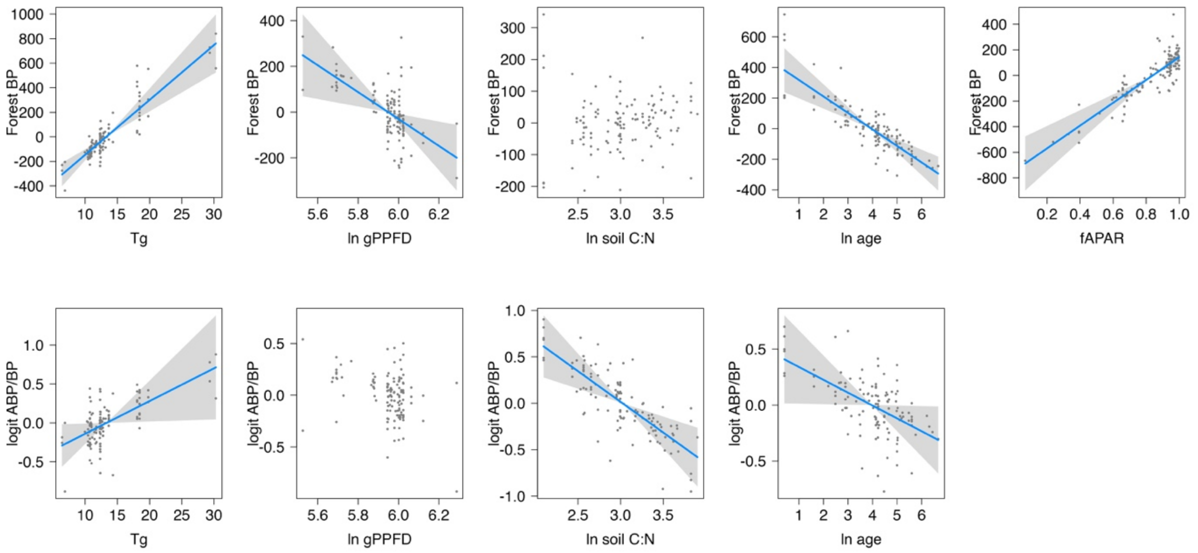


Fig. S5 Partial residual plots for statistical models developed to predict: (1) biomass production (BP; gC $m^{-2}yr^{-1}$) in forest; (2) the ratio of aboveground biomass production (ABP; gC $m^{-2}yr^{-1}$) to BP. Predictors are *measured* soil C/N, stand-age, fraction of absorbed photosynthetically active radiation (fAPAR), and incident photosynthetic photon flux density averaged over the growing season (gPPFD) and growth temperature ($T_g$). Predictors for soil C/N, age, gPPFD and $D$ were log-transformed. Statistical models used linear mixed-effects model, where site is a random intercept, and each point represents a measured value at ecosystem level.

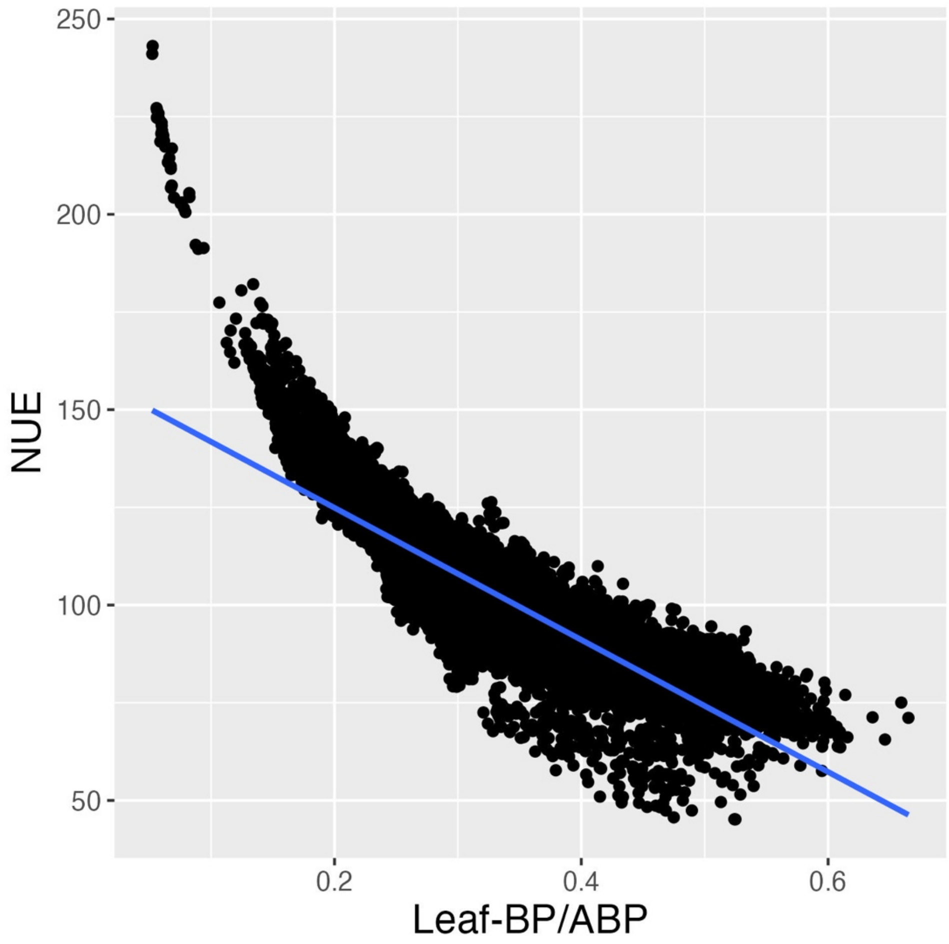


Fig. S6 Bi-variate relationship between nitrogen-use-efficiency (NUE) and leaf-BP/ABP basing on global prediction from data-driven model.
